# Effects of individuation and grouping on face representations in the visual cortex

**DOI:** 10.1101/423749

**Authors:** Hyehyeon Kim, Gayoung Kim, Sue-Hyun Lee

## Abstract

Top-down signals can influence our visual perception by providing guidance on information processing. Especially, top-down control between two basic frameworks, “Individuation” and “grouping”, is critical for information processing during face perception. Individuation of faces supports identity recognition while grouping subserves higher category level face perception such as race or gender. However, it still remains elusive how top-down dependent control between individuation and grouping affects cortical representations during face perception. Here we performed an fMRI experiment to investigate whether representations across early and high-level visual areas can be altered by top-down control between individuation and grouping process during face perception. Focusing on neural response patterns across the early visual cortex (EVC) and the face-selective area (the fusiform face area (FFA)), we found that the discriminability of individual faces from the response patterns was strong in the FFA but weak in the EVC during the individuation task whereas the EVC but not the FFA showed significant face discrimination during the grouping tasks. Thus, these findings suggest that the representation of face information across the early and high-level visual cortex is flexible depending on the top-down control of the perceptual framework between individuation and grouping.

## Introduction

Our visual perception depends on top-down signals such as attention or behavioral goals as well as bottom-up input (Gilbert and Li 2013). The brain is thought to efficiently use processing resources by selecting important information that is relevant to the current behavioral goal (Gilbert and Li 2013; Corradi-Dell’Acqua, Fink, and Weidner 2015). Top-down control of attention enables us to select behaviorally relevant stimuli from competing distracters (spatial attention) (Desimone and Duncan 1995), or focus on the relevant components (e.g., color or orientation)(feature attention) of a stimulus (Treue and Martinez Trujillo 1999; Motter 1994; Chelazzi et al. 1993). In another words, top-down signals can influence our visual perception by providing guidance or a framework for the information processing (Lupyan 2008; Goldstone and Hendrickson 2009; Goldstone, Lippa, and Shiffrin 2001). For example, when we see a chair, we usually identify it as a chair, but sometimes, we recognize it as furniture.

Especially, top-down control between two basic frameworks, “individuation” and “grouping”, is critical for the information processing during face perception. Individuation of faces supports identity recognition while grouping subserves higher category level face perception such as race or gender. Depending on the contexts, our face perception of others is based on a unique entity (individuation) or groups to which they belong such as race or gender (grouping) (Fiske and Neuberg 1990; Brewer 1988; Mason and Macrae 2004). For example, perceivers preferentially attend to the race category information of cross-race faces while they usually focus on the individuating information of same-race faces (Levin 2000; MacLin and Malpass 2001). It is generally thought that face individuation is preferentially done for in-group members, but grouping is done for out-group members (Levin 2000; Brewer 1988; Fiske and Neuberg 1990; Hugenberg et al. 2010; MacLin and Malpass 2001). Moreover, this preferential processing of individuation or grouping can be modulated by top-down task demands. Intentional individuation can contribute to better recognition of faces from other races (Hugenberg, Miller, and Claypool 2007; Pauker and Ambady 2009). In addition, DeLozier and Rhodes also showed that the value-directed motivation changed the own-race bias (DeLozier and Rhodes 2015).

The demands of top-down tasks affect the response patterns of the visual cortex during perception (Bugatus, Weiner, and Grill-Spector 2017; Nastase et al. 2017; Erez and Duncan 2015; Harel, Kravitz, and Baker 2014). Moreover, there is also evidence supporting that the activity of face-selective visual areas can be altered by different tasks (Wojciulik, Kanwisher, and Driver 1998; Cohen Kadosh et al. 2010; Reddy, Moradi, and Koch 2007; Gratton et al. 2013). Furthermore, recent studies have shown that the decoding of individual faces from the activity pattern of the fusiform face area (FFA) depends on the familiarity of faces (Hasinski and Sederberg 2016) or attended features of the faces (Anzellotti and Caramazza 2016; Dobs et al. 2018). These prior works suggest a possibility that top-down signals alter visual cortical representations. However, it remains still unclear how top-down control of individuation or grouping affects cortical representations during face perception.

Here we performed an fMRI experiment to investigate whether representations across the visual areas related to face processing can be altered by top-down control of individuation. Focusing on neural representations across the EVC and FFA, we found that the discrimination of individual faces is possible in the FFA but not in the EVC during the individuation task whereas the EVC but not the FFA showed a distinct neural activation pattern for each face during the grouping tasks. These results suggest that top-down control of the perceptual framework between individuation and grouping affects the representation of face information across the early and high-level visual cortex.

## Materials and Methods

### Participants

15 neurologically intact Korean participants (6 females; mean age = 23.67 ± 1.23 years) took part in the fMRI experiment (1 additional participant was excluded because of low performance in the individuation task which was less than chance level). 14 of the participants were right-handed and 1 participant was ambidextrous. All the participants provided a written informed consent for the procedure in accordance with the protocols approved by the KAIST Institutional Review Board.

### Stimuli

Participants were asked to perform individuation task, race grouping task, and gender grouping task in separate runs on the same set of sample faces. 8 sample face images were used for the sample phase of all tasks (**Figure 1**). The sample face images consisted of front-view photographs of 2 African-American men, 2 African-American women, 2 Caucasian men, and 2 Caucasian women, modified from the NimStim Face Stimulus Set (https://www.macbrain.org/resources.htm) (Tottenham et al. 2009) and Lifespan database of adult facial stimuli (http://agingmind.utdallas.edu/download-stimuli/face-database/) (Minear and Park 2004). During the test phase of the race or gender grouping task, a whole face image was presented. For this, 128 face images were selected and modified from the Chicago Face Database 2.0 (http://faculty.chicagobooth.edu/bernd.wittenbrink/cfd/index.html) (Ma, Correll, and Wittenbrink 2015). Every trial of the race or gender grouping task contained a new test image, which was never presented as a sample image and never reused in the other trials. During the test phase of the individuation task, a quarter-fragment of a face image either from the image presented during the sample phase of the trial or from the other sample image in its race and gender category was shown. Each sample and test image subtending approximately 7° of the visual angle was viewed at the center of the screen by a back-projection display (1024 × 768 resolution, 60 Hz refresh rate) with a uniform gray background.

**Figure 1.**
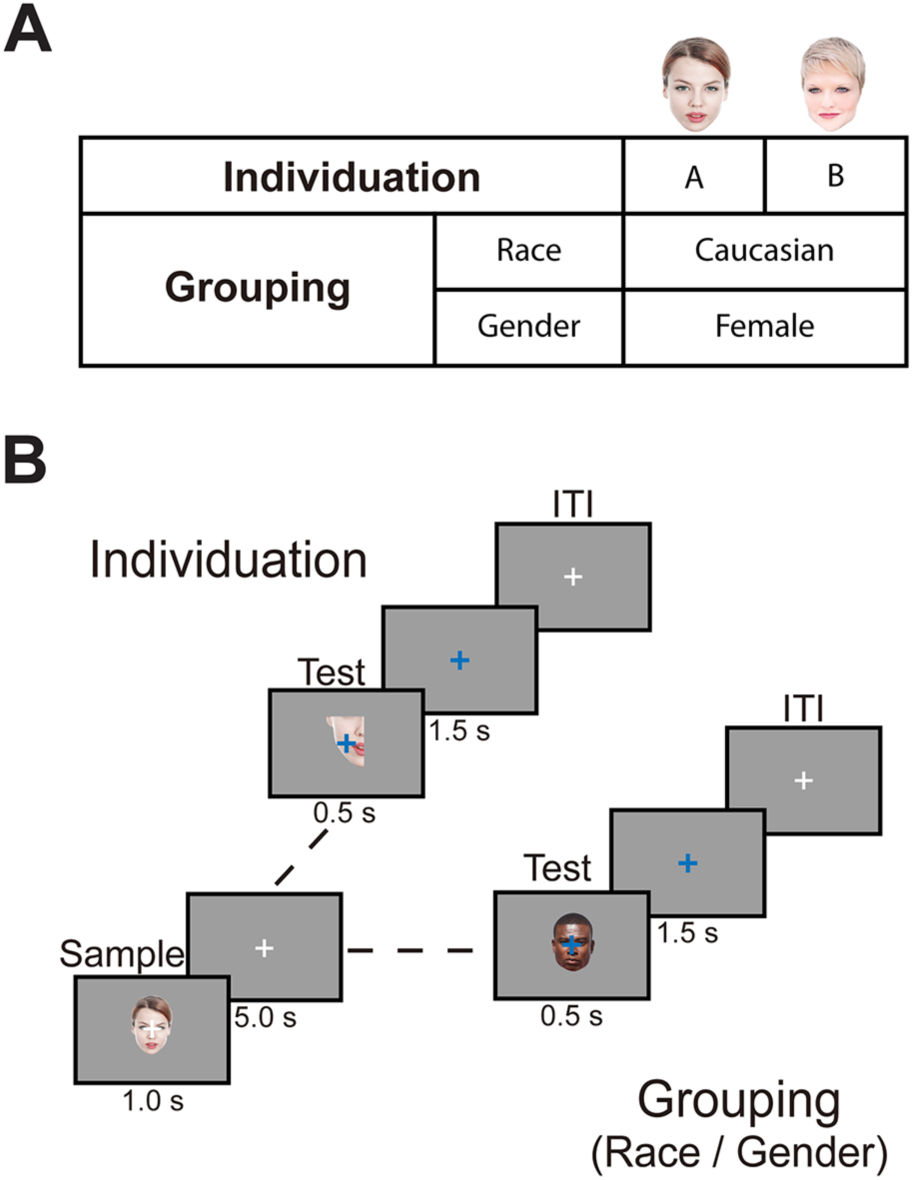
Experimental design. (A) Individuation and grouping frameworks for two sample face examples. In the individuation task, each face was perceived as unique entity whereas the groups (race or gender) to which the faces belong were emphasized in the grouping tasks. (B) One individuation task and two grouping tasks were performed in separate runs. On each trial for all tasks, a sample face from the same set was presented, followed by a 5-s interval and a test image. In the individuation task, the participants were instructed to determine whether each sample and the following test image was from the same face. In both of the grouping tasks, the participants were required to judge whether the sample and the test faces came from the same race (race grouping task) or gender (gender grouping task) category. The inter-trial interval (ITI) was randomized between 4 and 18 s. The fMRI analyses focused on the onset of each sample face presentation. The sample faces and test images were modified images from the NimStim Face Stimulus Set, Lifespan database, and Chicago Face Database 2.0, but in this Figure 1, the face images from the Pexels (www.pexels.com), licensed under the Creative Commons Zero (CC0) license, are used for illustration purpose only.

### Experimental Design

The participants performed three specific face perception tasks (one individuation task and two grouping (race and gender) tasks) during the event-related fMRI experiment. Each scan run contained one task, and the participants were instructed which task they would perform before each run. Each task was presented in 4 scan runs consisting of 16 trials each. The order of the runs was pseudorandomized to alternate the tasks. The order of the tasks was counterbalanced across the participants. The participants were asked to maintain fixation on a central cross throughout each run. No specific strategy for performing each task was provided to the participants.

On each trial of all tasks, a participant first saw a sample face image (sample phase), followed by a test image (test phase) (**Figure 1**). In the individuation task, the test image was a quarter-fragment of a face image, and the participants were asked to judge whether the test fragment belonged to the previously presented sample face image. They were instructed to press the ‘yes’ button if test image belonged to the sample face or the ‘no’ button if not as quickly as possible.

In both of the grouping tasks, the test image was a whole face image. The participants were required to judge whether the test face had the same common feature as the presented sample face. In the race grouping task, they needed to press the ‘yes’ button if the test image was from the same race category as the presented sample image or otherwise, press the ‘no’ button as quickly as possible. In the gender grouping task, they were asked to press the ‘yes’ button for the same gender test image as the sample face and the ‘no’ button’ for a different one.

On each trial of every task, a sample face was presented for 1 s, followed by a 5-s interval and a 500-ms test phase (**Figure 1**). The responses of the participants were recorded for 2 s from the beginning of the test image. The length of each trial was 8 s, and the inter-trial interval (ITI) varied with an average ITI of 6s. The order of the sample faces was randomized and counterbalanced across the runs. In a run, the number of trials for which the answer was “yes” (yes trial) and the trials for which the answer was “no” (no trial) were the same, and the order of the trials was randomized.

### Regions-of-interest (ROIs)

Two independent functional localizer scans were also collected from each participant to identify face-selective ROIs, including the FFA, and OFA. Each of these scans was an on/off block design with alternating 16-s blocks of different types of stimuli presented while the participants performed a one-back task. The participants were asked to maintain fixation on a central cross throughout each task. In the face localizer scan, 8 alternating blocks of face or object images subtending 7 degrees were presented in the center of the screen, and the first block was a face block. A total of 40 face images and 40 object images were used, and none of these face images overlapped with the face images used in the individuation, race, and gender grouping tasks. Thus, the face-selective ROIs were localized with the contrast of faces versus objects (Chan et al. 2010). ROIs that contained more than 15 voxels were included in the analysis. In one participant, the left OFA was not successfully localized, and another participant did not show right OFA. Thus, the BOLD responses of 14 participants were used for the left OFA or the right OFA analyses.

The retinotopic early visual cortex (EVC) (including V1, V2, V3, and V4) corresponding to the central visual field was defined by the localizer scan performed in independent set of participants; 14 participants viewed alternating 16s blocks of a central disk (5°) and an annulus (6-28°). The central EVC of each participant was transferred to the standard-mesh surface reconstructed by AFNI and SUMA (Saad and Reynolds 2012), and the overlap area across more than 4 participants was defined as the standard EVC. This standard EVC was re-transferred from the standard space to the space of each participant who performed the current face tasks. Voxels overlapped with the localized OFA were excluded for the analyses.

### fMRI Data Acquisition

Participants were scanned on the 3T Siemens MAGNETOM Prisma located in the Center for Neuroscience Imaging Research at the Institute for Basic Science. Echo-planar imaging (EPI) data were acquired using a 20 channel head coil with an in-plane resolution of 2 × 2 mm, and 35 2-mm slices (0.2-mm inter-slice gap, repetition time (TR) = 2 s, echo time (TE) = 25 ms, matrix size = 96 x 96, field of view (FOV) = 192 mm, and flip angle = 90°). Partial volumes of the temporal and occipital cortices were scanned, and our slices were oriented approximately parallel to the base of the temporal lobe. The order of all functional localizers and main task runs were pseudorandomized. Standard MPRAGE (magnetization-prepared rapid-acquisition gradient echo) images were collected with an in-plane resolution of 1 × 1 mm, and 1-mm slices (TR = 2.3 s, TE = 2.28 ms, FOV = 256 mm, flip angle = 8°) at the end of the experimental runs for each participant.

### fMRI Data Analysis

Data analysis was conducted using AFNI (http://afni.nimh.nih.gov) (Cox 1996), SUMA (AFNI surface mapper), FreeSurfer and custom MATLAB scripts. Data preprocessing included slice-time correction, motion correction, and smoothing (smoothing was used only for the localizer data with Gaussian blur of 5 mm full-width half-maximum (FWHM)). Then, the data were normalized by calculating the percent signal change for each subject on a voxel-by-voxel basis.

To derive the BOLD response magnitudes during the individuation, race, and gender grouping tasks, we conducted a standard general linear model using the AFNI software package (3dDeconvolve using GAM function) to deconvolve the event-related responses. The percentage BOLD signal change value and *t*-value of each voxel were derived from the onset of each sample phase.

For the average magnitude of responses, we calculated the average percentage BOLD signal change across all voxels and stimuli within each ROI. To derive the discrimination indices of the individual faces, we used the split-half correlation analysis method as the standard measure of information (Kravitz, Kriegeskorte, and Baker 2010; Haxby et al. 2001; Lee, Kravitz, and Baker 2012). For this, we first divided the 4 event-related runs for each task and each participant into 2 halves (each containing 2 runs) for all 3 possible ways. For each of the splits, we estimated the t-value between each event and baseline in each half of the data. The t-values were then extracted from the voxels within each ROI and cross-correlated. Here, the t-values were used rather than β-values because t-values tend to be more stable (Misaki et al. 2010), though we found nearly identical results from the analysis with the percentage BOLD signal change value. Before calculating the correlations, the *t*-values were normalized for each voxel by subtracting the mean value across all face conditions (Lee, Kravitz, and Baker 2012; Haxby et al. 2001). For each split, the Pearson correlation coefficients between all possible pairs of activation patterns for 8 stimuli from 2 halves of the data were calculated and Fisher-z-transformed. Then, the within-correlations were derived by calculating the average correlation between the same face conditions from each split, whereas the between-correlation was calculated as an average correlation between the different face conditions from each split. A discrimination index for a face condition was defined by subtracting the average of the between-condition correlations from the within-condition correlations.

ROIs were created for each participant from the localizer scans. Significance maps of the brain were computed by performing a correlation analysis thresholded at a p value of 0.0001 (uncorrected). ROIs were generated from these maps by taking the contiguous clusters of the voxels that exceeded the threshold and occupied the appropriate anatomical location based on previous studies (Sayres and Grill-Spector 2008; Schwarzlose et al. 2008).

### Statistical Analyses

To compare within-correlations with between-correlations, we used one-sample t-tests (one-tailed) with the assumption that within-correlations are greater than between-correlations. To compare discrimination indices with basal level (zero), one-sample t-tests (one-tailed) were used with the assumption of a predicted positive direction. Repeated-measures ANOVAs (tests of within-subjects effects) were used to determine the statistical significance of a task or the ROI effects. For all ANOVAs with factors with more than two levels, Greenhouse-Geisser Corrections were used.

## Results

To investigate how top-down control of individuation affects neural responses during face perception, we asked participants to perform one individuation task and two grouping tasks in separate runs on the same set of eight different faces. The tasks were designed to emphasize the individual unique characteristics of faces (individuation task) or categorization according to a common feature such as race (race grouping task) or gender (gender grouping task) during perception (**Figure 1**). In each trial for the tasks, the participants were asked to see one of the sample faces, followed by a test image. The individuation task was designed to emphasize faces on an individual level during the sample face perception. In this task, the participants had to determine whether each sample and following test image were from the same face or not. The grouping tasks were designed to emphasize race (race grouping task) or gender (gender grouping task) category during the sample face perception. In these tasks, the participants were asked to determine whether the sample and the test faces came from the same race or gender category. Task performance accuracy was greater than the chance levels in all tasks (t(14) = 12.19, p = 7.63 × 10^-9^ for the individuation task; t(14) = 24.71, p = 6.02 × 10^-13^ for the race grouping task; t(14) = 36.55, p = 2.72 × 10^-15^ for the gender grouping task) (**Supplementary Table 1**), indicating that the participants successfully performed each task.

### Average activation

Because the FFA and OFA have been considered as the high-level visual areas that are associated with face perception (Kanwisher, McDermott, and Chun 1997; Rhodes et al. 2009; Haxby, Hoffman, and Gobbini 2000), we focused on the neural responses of the FFA and OFA as well as the EVC (**Figure 2A**). We first performed univariate analysis (**Supplementary Figure 1**). We derived the average BOLD signal changes for the onset of each sample phase. The average BOLD signal changes were significantly positive in all ROIs across all tasks (FFA, t(14) > 2.80, p < 0.02; OFA, t(13) > 2.52, p < 0.03; EVC, t(14) > 3.96, p < 0.01), except for the signal change of the left OFA during the gender grouping task (t(13) = 1.87, p = 0.08). A three-way ANOVA with Task (individuation, race, gender), ROI (FFA, OFA, EVC) and hemisphere (left, right) as within-subject factors revealed a highly significant main effect of Task (F(1.49, 17.84) = 7.28, p = 0.01) as well as a main effect of ROI (F(1.36, 16.35) = 5.35, p = 0.03) but no main effects of hemisphere (F(1,12) = 0.53, p = 0.48). These results suggest that the average neural responses of the FFA, OFA, and EVC during perception vary depending on tasks even for the same set of faces.

**Figure 2.**
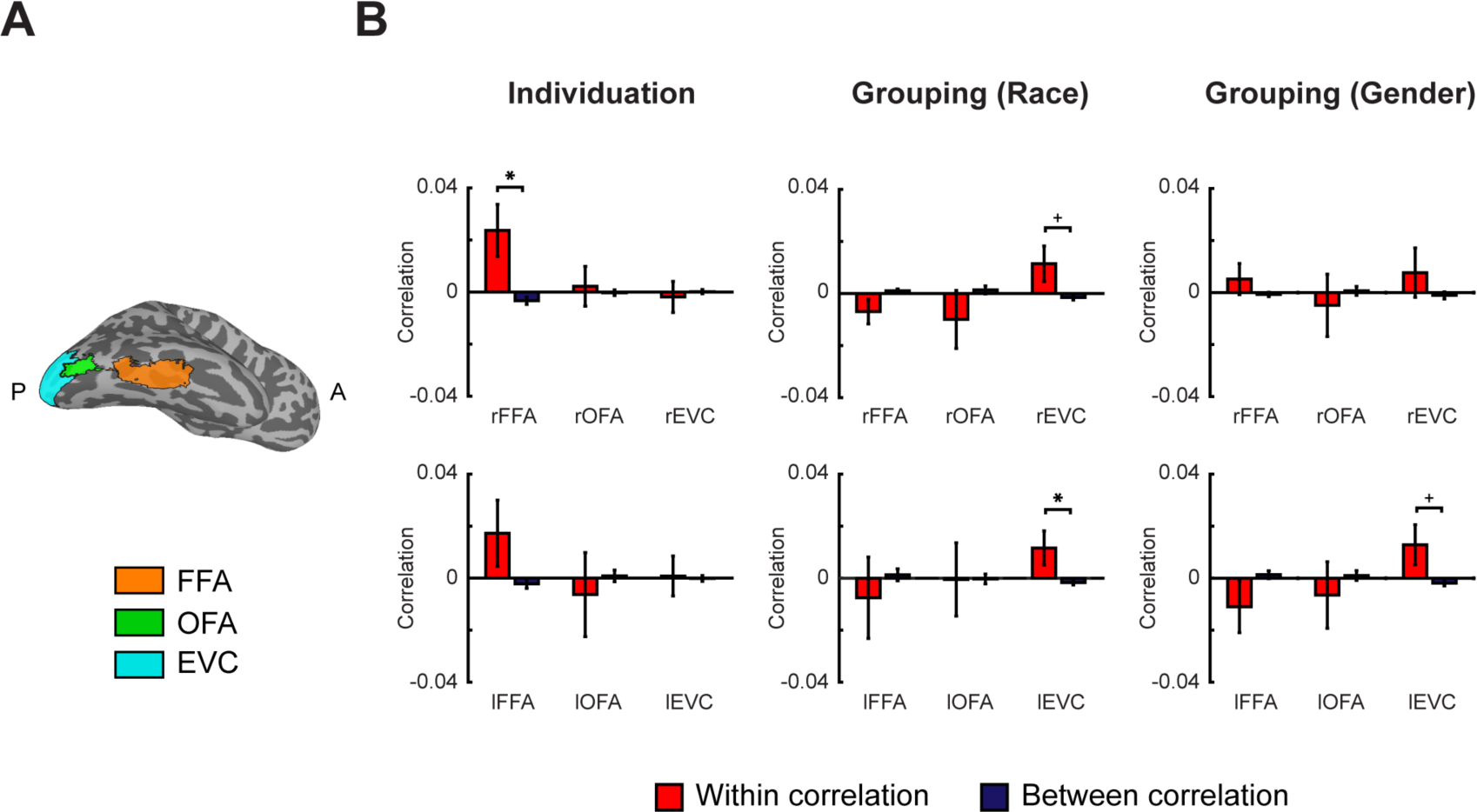
Individual face discrimination from the neural responses of the FF A, OF A, and EVC ROIs in each hemisphere during the individuation and grouping (race and gender) tasks. (A) Regions-of-interest (ROIs). For illustrative purposes, ROIs of the right hemisphere derived from each participant were transformed to a standard space. Orange (FFA), green (OFA), or blue (EVC) areas indicate voxels which were with the ROI in at least 4 participants. A, Anterior; P, Posterior (B) During the individuation task, significantly greater within-correlations than between-correlations were found in the right FFA (rFFA) but not in the other ROIs whereas the left EVC but not the other ROIs showed greater within-correlation values than between-correlation values during the race or gender task. ^∗^p < 0.05. +p = 0.05. Error bars indicated between-subjects s.e.m.

### Individual face information across the visual cortex

To directly investigate the neural representations of individual faces, we next used multivariate pattern analysis (MVPA). For this, we compared the patterns of responses across two independent halves of data, which were extracted within each ROI. The within-correlation values (correlation within the same face conditions) and the between-correlation values (correlation between different face conditions) were obtained for all combinations of sets. A higher within-correlation than a between-correlation for a ROI indicates that the ROI shows individual face specific response patterns (Haxby et al. 2001; Lee, Kravitz, and Baker 2012) (See **Materials and Methods** for details). During the individuation task, significantly greater within-correlations than between-correlations were found in the right FFA (rFFA, t(14) = 2.45, p = 0.01) but not in other visual areas (rOFA, t(13) = 0.28, p = 0.39; rEVC, t(14) = −0.31, p = 0.62; lFFA, t(14) = 1.40, p = 0.09; lOFA, t(13) = −0.39, p = 0.65; lEVC, t(14) = 0.11, p = 0.46) (**Figure 2B**). Thus, this suggests that the discriminability of individual faces from the response patterns was strong in the rFFA but weak in other visual cortical areas during the individuation task.

During the race grouping task, the left EVC (lEVC, t(14) = 1.84, p = 0.04) and the right EVC (rEVC, t(14) = 1.72, p = 0.05) but not other visual areas (rFFA, t(14) = −1.53, p = 0.93; rOFA, t(13) = −0.89, p = 0.81; lFFA, t(14) = −0.51, p = 0.69; lOFA, t(13) = −0.01, p = 0.51) showed significantly greater within-correlation values than between-correlation values (**Figure 2B**). Moreover, the same tendency of within- and between correlations was also found during the gender grouping task (**Figure 2B**). For the within-correlation values of the race and gender grouping tasks, a three-way ANOVA with Task (race and gender grouping tasks), ROI (FFA, OFA, EVC) and hemisphere (right, left) as within-subject factors revealed the main effect of ROI (F(1.86, 22.32) = 6.17, p = 0.01) but no effect of task (F(1,12) = 0.09, p = 0.77) or hemisphere (F(1,12) = 0.01, p = 0.92). Thus, this suggests that the distribution of individual face information across the FFA, OFA and EVC is similar in both grouping tasks, and that there is no statistically significant difference between the left and right hemisphere.

We observed individual face discrimination in the face-selective area (FFA) but not in the EVC during the individuation task, whereas the opposite was true during grouping tasks (**Figure 2**). To further examine this property, we derived the discrimination indices by subtracting the between-correlation values from the within-correlation values (**Figure 3**). Given that there were similar profiles for the within- and between correlations across the ROIs between the right and left hemispheres (**Figure 2B**), we collapsed across the hemispheres. To directly compare the shared responses in both grouping tasks with the responses of the individuation task, we derived average response patterns across the grouping tasks and directly compared the discrimination indices between the individuation and grouping tasks (**Figure 3**). Consistent with the previous analysis data, during the individuation task, the FFA (t(14) = 2.10, p = 0.03) but not the EVC (t(14) = −0.12, p = 0.55) showed significant discrimination for individual faces, whereas during the grouping tasks, the opposite pattern was observed, with significant decoding of faces in the EVC (t(14) = 3.31, p < 0.01) but not in the FFA (t(14) = −0.55, p = 0.70). A two-way ANOVA with Task (individuation and grouping tasks) and ROI (FFA and EVC) as the within-subject factors showed a significant interaction between the task and ROI (F(1,14) = 24.91, p < 0.01), reflecting the task-dependent dissociation of the FFA and EVC; a strong face decoding in the FFA and weak decoding in the EVC during the individuation task, but the opposite pattern during the grouping task. Thus, while multiple regions are activated by face perception, they flexibly represent face information depending on the top-down control of the perceptual framework between individuation and grouping.

**Figure 3.**
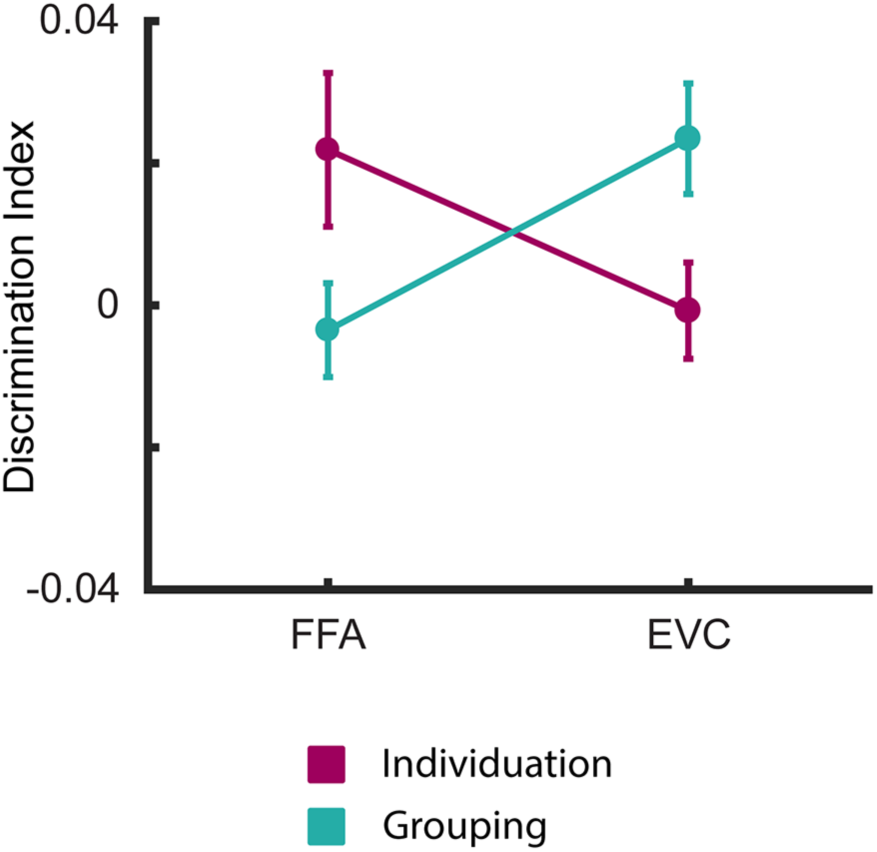
Task-dependent dissociation of the FFA and EVC. The FFA but not the EVC showed significant discrimination of individual faces during the individuation task, whereas the opposite pattern was observed during the grouping tasks. Error bars indicated between-subjects s.e.m.

## Discussion

Our findings demonstrate distinct neural representations for faces in individuation and grouping conditions. Using multivoxel pattern analysis, we found that cortical representations for individual faces were strong in the high-level face-selective area (FFA) but weak in the EVC during the individuation task whereas the EVC but not the FFA contained individual face information during the grouping tasks. Thus, these suggest that visual cortical regions are flexibly recruited for representations of individual faces depending on the top-down signals to control the perceptual framework between individuation and grouping.

Prior studies have shown hierarchical visual representations (Horikawa and Kamitani 2017; Riesenhuber and Poggio 1999; DiCarlo, Zoccolan, and Rust 2012). Especially for face perception, it has been suggested that the EVC is mainly involved in the representation of lower-order visual features of faces such as skin color (Brosch, Bar-David, and Phelps 2013) or forehead size (Kaul, Rees, and Ishai 2011) while neural responses of the FFA more reflect higher-order features such as face identity (Haxby, Hoffman, and Gobbini 2000; Grill-Spector, Knouf, and Kanwisher 2004; Hoffman and Haxby 2000; Rotshtein et al. 2005; Winston et al. 2004; Gauthier et al. 2000). The OFA shows intermediate properties of representations between the EVC and the FFA (Ramon, Dricot, and Rossion 2010; Rossion et al. 2003; Rotshtein et al. 2005; Haxby, Hoffman, and Gobbini 2000). Recently, Guntupalli et al. suggested a progressive disentangling of the representation of face identity from the representation of head view, showing that while the early visual cortex and the OFA distinguish head view but not identity, representation of identities was achieved in the FFA (Guntupalli, Wheeler, and Gobbini 2017). Based on these prior works, the flexible recruitment of cortical regions for individual face representations in our data may reflect top-down dependent emphasis of hierarchically different features; for individuation, high-order features may be critical, while low-order features can be mainly recruited for a grouping condition. This top-down control dependent dissociation of the FFA and EVC suggests a possible explanation for why face identity decoding was not successful in some studies while it was possible in others in the face-selective areas (Anzellotti, Fairhall, and Caramazza 2014; Axelrod and Yovel 2015; Kriegeskorte et al. 2007; Hasinski and Sederberg 2016; Guntupalli, Wheeler, and Gobbini 2017).

Beyond the “core” regions of face perception such as the FFA and the OFA, it is known that extended brain regions including the anterior temporal lobe area, amygdala, orbitofrontal cortex, and the inferior frontal gyrus also collectively contribute to face perception (Phillips et al. 1997; Leveroni et al. 2000; Ishai., Haxby, and Ungerleider 2002; O’Doherty et al. 2003; Kaul, Rees, and Ishai 2011; Axelrod and Yovel 2015; Kriegeskorte et al. 2007; Anzellotti, Fairhall, and Caramazza 2014). Especially, the anterior temporal lobe is thought to be engaged in face identification (Kriegeskorte et al. 2007; Anzellotti, Fairhall, and Caramazza 2014; Axelrod and Yovel 2015). The inferior frontal gyrus is considered to process semantic aspects of faces (Leveroni et al. 2000; Ishai., Haxby, and Ungerleider 2002; Kaul, Rees, and Ishai 2011), and therefore it is likely to have a role in connecting the current behavioral goal with the face perception process. It will be interesting to further investigate the role of extended face-selective regions and their response changes according to top-down signals.

In the present study, we focused on the representation of individual face information in individuation and grouping conditions. We additionally examined the representations of race or gender information across the EVC, OFA, and FFA, but could not find the opposite gradient of information across the ROIs between the individuation and grouping conditions, which was found in the representation of individual face information. Given a recent study in which social-conceptual knowledge altered the representations of gender, race, and emotion categories (Stolier and Freeman 2016), the effect of subjective memories or experiences might influence more broad areas such as the prefrontal cortex rather than being restricted to the visual cortex. Additionally, our data do not totally exclude the possibility that task difficulty or general arousal contributes to the distinction of the information gradient between the individuation and grouping conditions. Future studies are needed to investigate these possibilities.

In summary, our results show that a dissociation of the FFA and EVC for individual face representations in individuation and grouping conditions; there is significant individual face information in the FFA but weak information in the EVC during individuation whereas the opposite pattern is found during grouping. These results provide evidence for that the visual representations during face perception are flexible depending on the top-down control of the individuation process.

## Acknowledgements

This work was supported by a grant of the Korea Health Technology R&D Project through the Korea Health Industry Development Institute (KHIDI) funded by the Ministry of Health & Welfare (HI15C3175), and the Basic Science Research Program NRF-2016R1C1B2010726) and the Brain Research Program (NRF-2017M3C7A1031333) through the National Research Foundation (NRF) of Korea. Thanks to members of the Memory and Cognition Laboratory, KAIST for discussion.

## Author Contributions

S.-H.L. designed and supervised the research. H.K. performed the research, and H.K. and G.K. analyzed the data. H.K., G.K. and S.-H.L. wrote the manuscript.

